# Estimate of the mutation rate in the endangered Devils Hole pupfish provides equivocal support for the drift-barrier hypothesis at an outlying extreme

**DOI:** 10.64898/2026.01.11.698928

**Authors:** David Tian, Elise Kerdoncuff, Kevin Wilson, Olin Feuerbacher, Michael R. Schwemm, Jennifer Gumm, Priya Moorjani, Christopher H. Martin

## Abstract

Mutation rates vary by orders of magnitude across eukaryotes. The drift-barrier hypothesis proposes that drift overwhelms selection for lower mutation rates in small populations, leading to higher mutation rates over time due to the gradual accumulation of mutator alleles. Here, we test this hypothesis in one of the smallest long-term isolated populations in the world, the endangered Devils Hole pupfish (*Cyprinodon diabolis*). We estimated germline mutation rates in adults and embryos that died prematurely using autozygous segments due to recent inbreeding events. Our estimate of 8.09 × 10^−9^ per base pair per generation was higher than the mean rate for actinopterygian fishes of 5.97 × 10^−9^ (95% CI: 4.39 × 10^−9^ - 7.55 × 10^−9^) but was lower than predicted for such a low effective population size based on a recent meta-analysis of vertebrate mutation rates and similar to species with much larger effective population sizes, contradicting the drift barrier hypothesis. We also found that embryos which died during development had a higher mutation rate than mature adults, potentially reflecting a segregating lethal mutator allele. We analyzed the mutational spectra of germline mutations and found that spectra between embryonic lethal and mature adults were similar and comparable to other fishes, despite environmental differences in temperature, oxygen saturation, and UV exposure. Mutation rates in this endangered species provide new insights into the mechanisms driving mutation rate variation across vertebrates at one extreme of low effective population size in nature.

## Introduction

Germline mutations provide the raw material for evolution (Barton and Keightley 2002). The rate of germline mutations varies by three orders of magnitude across multicellular eukaryotes (Baer et al. 2007; Wang and Obbard 2023) and this has many implications for selection, drift, mutation load, and the adaptive capacity of populations. In recent years, a range of pedigree-based mutation rate estimates have been published, revealing that per-generation mutation rates across vertebrates vary substantially by at least an order of magnitude (Feng et al. 2017; Chintalapati and Moorjani 2020; Wang et al. 2022; Bergeron et al. 2023; de Manuel et al. 2025; Peña-García et al. 2025). This variation has been attributed to myriad life-history traits, such as generation time (Ohta 1993; Bergeron et al. 2023; Wang and Obbard 2023), age at sexual maturity (Bergeron et al. 2023), fecundity (Bergeron et al. 2023), metabolic rate (Martin and Palumbi 1993), longevity (Wilson et al. 2011; Thomas et al. 2018; Zhu et al. 2025), mating strategy (Bergeron et al. 2023), and body size (Bergeron et al. 2023). Stressful environments (Liu and Zhang 2019; Wei et al. 2022), greater exposure to UV radiation (Hoeijmakers 2009), and oxidative stress (Cadet and Wagner 2013; Poetsch et al. 2018; Kessler et al. 2020) are also known to affect mutation rates. Molecular architecture also plays a role, including DNA damage and repair mechanisms (Gao et al. 2019; Volkova et al. 2020), DNA methylation (Wolfe et al. 1989; Mugal et al. 2015), replication timing (Stamatoyannopoulos et al. 2009), genome size (Wang and Obbard 2023), fragile structural arrangements (Xie et al. 2019), and microsatellite instability (Jeffreys et al. 1988).

An additional mechanism potentially driving mutation rate variation is the drift-barrier hypothesis: a negative correlation between effective population size and mutation rate due to purging of deleterious mutations that increase mutation rates in larger populations; whereas smaller populations have reduced efficacy of selection resulting in mutator alleles persisting and becoming fixed in the population over time due to drift and increasing mutation rates (Kimura 1967; Sung et al. 2012; Lynch et al. 2016; Bergeron et al. 2023; Lynch et al. 2023; de Manuel et al. 2025). This pattern originally appeared consistent across vertebrate taxa (Bergeron et al. 2023) and may possibly hold more broadly across eukaryotes and prokaryotes (Lynch 2010). However, population size and generation time are inversely correlated (Chao and Carr 1993; Waples et al. 2013) and mutation rates tend to be higher in species with longer generation times and lifespan (Bergeron et al. 2023; Wang and Obbard 2023). A recent reanalysis of the data in Bergeron et al. (2023) by Weinstein and Roy (2026) found that that once generation time was accounted for, there was no relationship between mutation rate and effective population size.

Nonetheless, few examples of species with consistently small long-term effective population sizes have been measured (but see Wayne et al. 1991 and Foote et al. 2021). To disentangle the impact of generation time and population size, studies of outlier species with a range of generation times and effective population sizes are necessary. In this study, we leverage the short generation time and long-term small effective population size of the Devils Hole pupfish (*Cyprinodon diabolis*), one of the most endangered species in the world, to test the drift-barrier hypothesis at its most extreme limits. The drift-barrier hypothesis predicts that the Devils Hole pupfish should have one of the highest mutation rates among vertebrates.

*Cyprinodon diabolis* is found only in Devils Hole, a 3.5 m x 22 m cavern that is the smallest vertebrate range in the world (Deacon et al. 1995). Genomic and phylogenetic estimates suggest that Devils Hole pupfish have been isolated in Devils Hole for several hundred to several thousand years (Martin et al. 2016; Martin et al. 2017). Devils Hole is an extremely inhospitable environment for fishes due to a constant water temperature of 33-34°C (Miller 1948; James 1969), only 9 months of direct sunlight per year which limits nutrient availability (Wilson et al. 2001; Riggs and Deacon 2002), and low dissolved oxygen levels near lethal limits for fishes (Gustafson and Deacon 1997). The species is also the smallest *Cyprinodon* species with the shortest lifespan and shortest generation time of just one year (Stoike and Pister 2010). Biannual population census counts since 1972 show that the population was initially relatively stable between around 200 and 500 individuals in the spring and fall (Tian et al. 2022). However, the population began to decline in the mid-1990s, with official counts showing lows of 38, 35, and 38 in 2007, 2013, and 2025 respectively. Further, unofficial counts show the population reached an all-time low of only 20 individuals in early 2025. Estimates of effective population size based on the harmonic mean of census population size range between 122 (spring) and 209 (fall) (Tian et al. 2022). Due to its long-term small effective population size, stressful food-limited environment, increased metabolic rate due to extremely high year-round temperatures, small body size, and short generation time, we hypothesized that Devils Hole pupfish may have one of the highest mutation rates among all vertebrates (Martin and Höhna 2018). Unfortunately, no pedigrees are available for this endangered species due to its restricted breeding at the U.S. Fish and Wildlife Service Ash Meadows Fish Conservation Facility. Consequently, we estimated the mutation rate by leveraging the high levels of inbreeding within Devils Hole pupfish (Tian et al. 2022) resulting in large numbers of autozygous segments within the genome, an approach that has successfully generated accurate mutation rate estimates in human populations (Campbell et al. 2012; Narasimhan et al. 2017).

## Results

### Sampling

We sampled 15 wild adults, 25 captive adults, and 22 embryonic lethal individuals, embryos that died during development at 5-7 days post fertilization (dpf), for a total of 62 individuals. Sequencing of adults was based on caudal fin clips of adult fish reared in the Ash Meadows Fish Conservation Facility derived from wild or refuge-collected eggs, while sequencing of embryonic lethals was based on embryos showing visual indicators of developmental slowdown and reduced heart rate at 5-7 dpf indicating impending death. These indicators were previously confirmed to be 100% lethal. 58 samples were sequenced to medium-coverage (mean coverage = 18X) while an additional 4 refuge adult samples were sequenced to high-coverage (mean coverage = 98X) for a total of 62 sequenced genomes (Supp. Table 1).

### Distribution of autozygous segments

We identified a total of 2,754 autozygous segments across 62 individuals using the hidden Markov model (HMM) introduced in Narasimhan (2016), including 526 segments that were at least 5 Mb in length across 42 individuals. The number of segments per individual varied (mean = 12.5, median = 8, range = 1 – 46). Following Narasimhan (2017), we trimmed the ends of each autozygous segment by removing 1MB from both sides to avoid uncertainty in the calling of autozygous breakpoints and accordingly adjusted the number of callable sites. After filtering, we retained 2.99 Gb across 42 individuals. Using the distribution of autozygous segment lengths across the genome, we inferred a weighted mean of 7.54 meioses (adults = 8.35, embryonic lethals = 7.19), weighted by the proportion of the genome in autozygous segments greater than 5 Mb (range per individual = 6.2 – 15.9) across our dataset (Fig. 1).

**Fig. 1.**
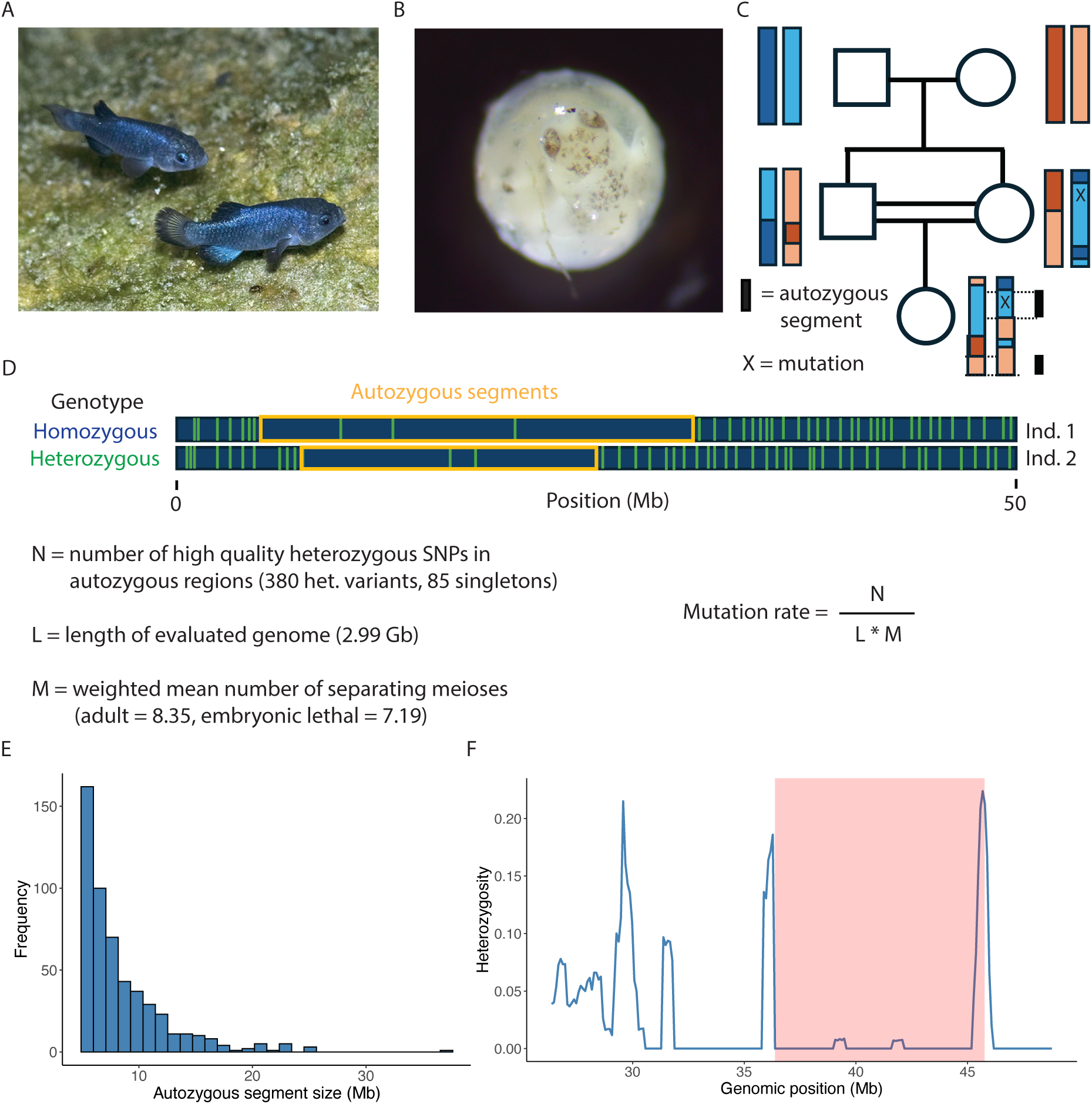
Mutation rate estimation. (a) *C. diabolis* within Devils Hole by Olin Feuerbacher. (b) Embryonic lethal *C. diabolis* that died during development 5-7 days post fertilization. (c) Example pedigree where inbreeding generates autozygous segments and the opportunity to identify recent mutations. (d) Heterozygous sites (green) in autozygous segments (yellow) are inferred to be recent mutations. Calculation of the mutation rate based on N, L, and M, as described in the methods. (e) Histogram of 526 identified autozygous segments at least 5 Mb long. (f) Sliding windows of heterozygosity and an inferred autozygous segment in shaded red in individual RT17, located at Chr6: 36375692-45758493. Note that calling autozygous segments involves some uncertainty around the cutoff points; thus, we excluded 1 Mb from both ends of each segment for our analyses.

### Mutation rate estimates

The 526 autozygous segments among 42 individuals contained 380 total heterozygous variants that are potential mutations, including 85 singletons (Supp. Table 2). The transition-transversion ratio among our heterozygous variants was 1.29, similar to that of sticklebacks (1.23) (Zhang et al. 2023) and Malawi cichlids (1.73) (Malinsky et al. 2018), suggesting that our identification of potential mutations are reliable. Several individuals had very few segments larger than 5 Mb and thus no mutations were detected; these individuals were excluded from our estimate.

We estimated the mutation rate in two different ways. First, we used the count of singleton heterozygotes found in autozygous regions (# singletons / length of callable bp * 2) to estimate the *de novo* mutation rate to be 8.09 × 10^−9^ per base pair per generation (95% CI: 4.20 × 10^−9^ – 1.27 × 10^−8^) among our adult individuals (Fig. 2). Our singleton-based estimates for our high-coverage (mean = 98X) and medium-coverage (mean = 18X) adult samples were similar at 8.28 × 10^−9^ per base pair per generation (95% CI: 4.45 × 10^−9^ – 1.28 × 10^−8^) and 7.45 × 10^−9^ per base pair per generation (95% CI: 0 – 1.87 × 10^−8^), respectively.

**Fig. 2.**
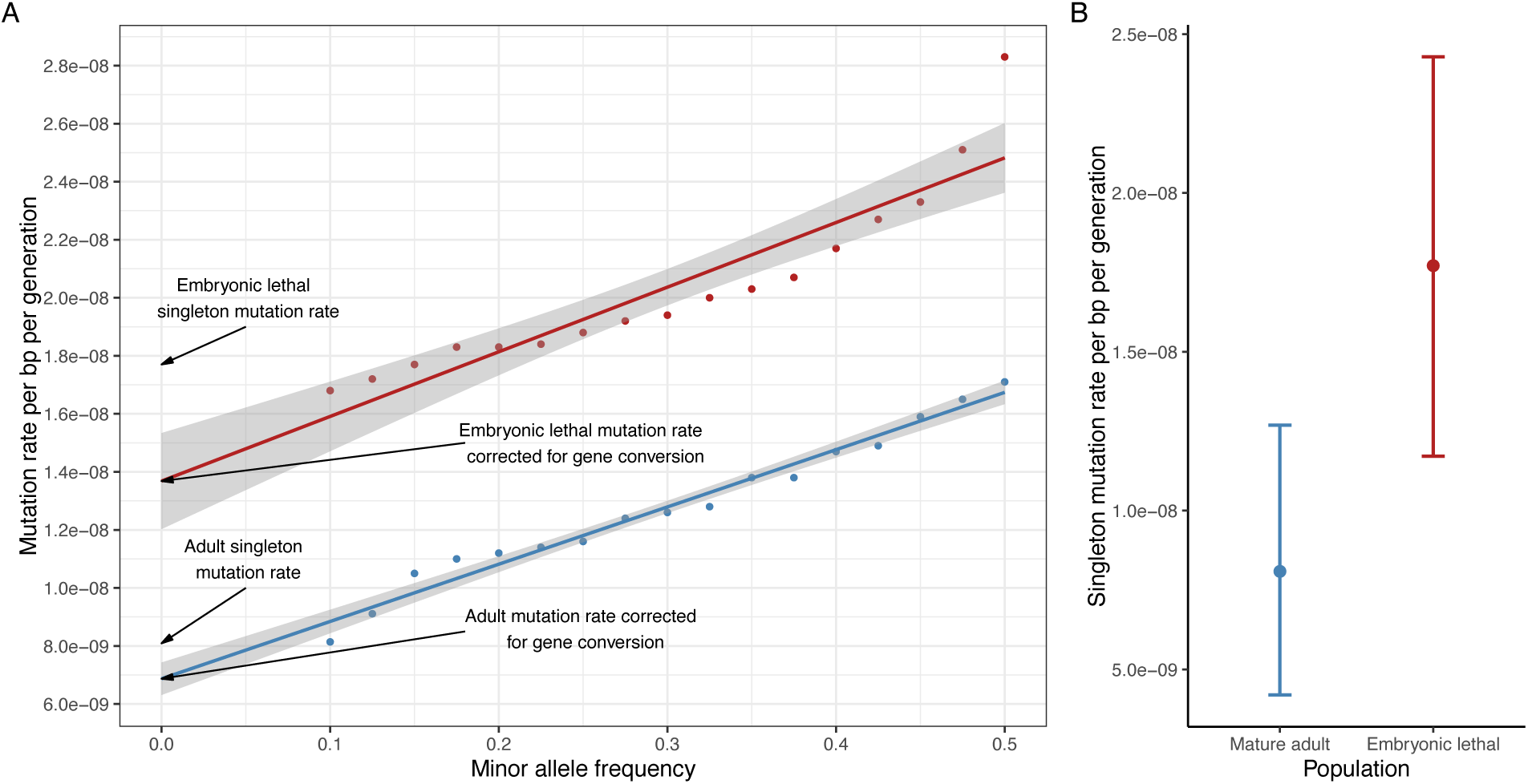
Estimate of the mutation rate in the Devils Hole pupfish. (a) Minor allele frequency threshold regression to estimate the mutation rate while accounting for gene conversion for both embryonic lethal (red) and mature adult (blue) individuals. The mutation rate is calculated by obtaining the number of mismatches in autozygous segments at different thresholds of minor allele frequency. The y-intercept represents the mutation rate corrected for gene conversion. (b) Weighted-mean singleton mutation rates weighted by the number of callable bp in autozygous regions greater than 5 Mb per individual. Error bars show bootstrap 95% confidence intervals.

Second, we used minor allele frequency threshold regression (Palamara et al. 2015) to estimate a mutation rate corrected for gene conversion (# heterozygotes below a given MAF threshold / length of callable bp * weighted average number of meioses). This approach resulted in an estimated mutation rate of 6.87 × 10^−9^ per base pair per generation (95% CI: 6.31 × 10^−9^– 7.43 × 10^−9^) and a non-crossover gene conversion rate of 7.41 × 10^−4^ per base pair per generation (Fig. 2). We note that the confidence intervals for our singleton and gene conversion corrected estimates of the adult mutation rate overlap. However, given the additional uncertainty surrounding the inferred number of meioses from a limited number of autozygous segments, we have higher confidence in our singleton mutation rate estimate, which does not rely on inference of meioses past a single generation.

Following the same approach for the embryonic lethal *C. diabolis* samples, we estimated a singleton mutation rate of 1.77 × 10^−8^ per base pair per generation (95% CI: 1.17 × 10^−8^ – 2.43 × 10^−8^) and found the difference between adult and embryonic lethal singleton mutation rates to be significantly different from zero (95% CI: 2.35 × 10^−9^ – 1.72 × 10^−8^, *P* = 0.0112). We estimated a gene conversion corrected mutation rate for embryonic lethal samples of 1.37 × 10^−8^ per base pair per generation (95% CI: 1.20 × 10^−8^– 1.53 × 10^−8^), once again overlapping with the confidence intervals of the singleton-based embryonic lethal mutation rate estimate.

### Estimates of effective population size

We used our estimated mutation rate to calculate effective population sizes in the wild and captive refuge populations. Paired with previously calculated measures of π (Tian et al. 2026), we estimated effective population size given the relationship of *N_e_* = π / 4µ to be 426. This is consistent with historical census sizes in wild populations of 200 – 500 individuals (Deacon 1979), and slightly higher than previous estimates of effective population size based on the harmonic mean of wild census population size over time (spring = 122, fall = 209, mean = 165) (Tian et al. 2022).

### Mutation rate in the context of life history traits and effective population size

Relative to other vertebrates, the adult *C. diabolis* mutation rate is on the upper end but consistent with expected mutation rates as a function of generation time, age at sexual maturity, and lifespan in the wild (Fig. 3). Our estimated mutation rate of 8.09 × 10^−9^ per base pair per generation for adults and 1.77 × 10^−8^ per base pair per generation for embryonic lethal individuals is low but within the range of what the drift-barrier hypothesis predicts for a species with an *N_e_* of 165 based on a phylogenetically-corrected linear regression across vertebrate taxa from Bergeron et al. (2023) that predicts a mutation rate of 1.23 × 10^−8^ per base pair per generation (95% CI = 7.81 × 10^−9^ – 1.93 × 10^−8^) (Fig. 3d) and provides missing data at the extreme end of existing mutation rate estimates. Our adult estimate is also low but within the range of the drift-barrier prediction of 1.12 × 10^−8^ per base pair per generation (95% CI = 7.63 × 10^−9^ – 1.64 × 10^−8^), when using an *N_e_* of 426, derived from *N_e_* = π / 4µ.

**Fig. 3.**
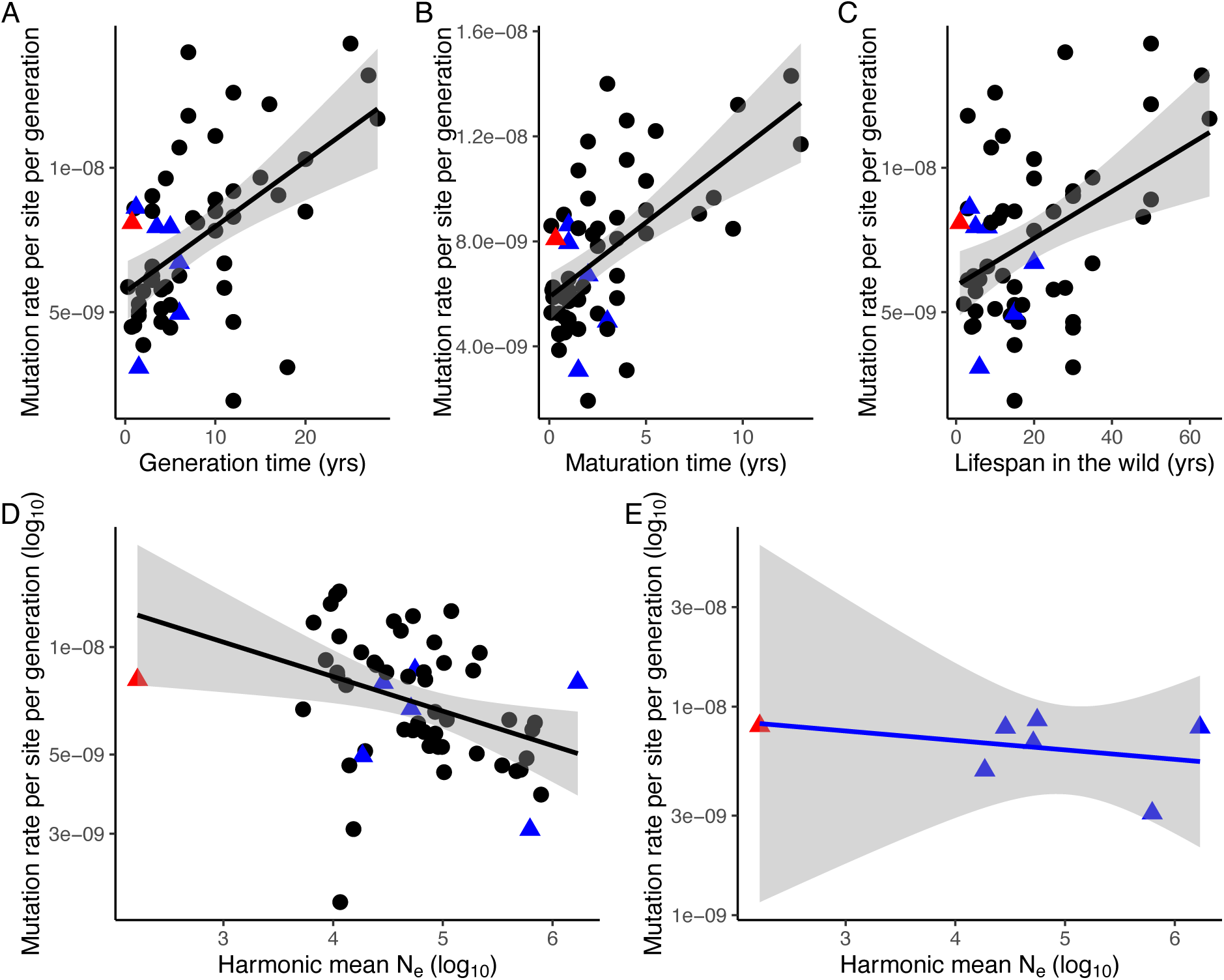
*C. diabolis* mutation rate relative to other vertebrate mutation rates. Mutation rates in the context of the following life-history traits: (a) generation time (9 months), (b) age at sexual maturity (4 months), (c) lifespan in the wild (1 year), (d) effective population size across vertebrates (*N_e_* = 165), and (e) effective population size across ray-finned fishes (*N_e_* = 165). Shading represents 95% confidence interval. The red triangle represents *C. diabolis*, while the blue triangles and black dots represent other ray-finned fishes and tetrapods, respectively, from Bergeron et al. 2023. For *C. diabolis*, we estimated *N_e_* as the average of the harmonic means of spring and fall census sizes from 1970 to 2020. In contrast, the *N_e_* in Bergeron et al. (2023) is based on the harmonic mean of population sizes estimated by PSMC between 30,000 and 1,000,000 years.

However, when restricted to the six pedigree mutation rate estimates with effective population sizes available for actinopterygian fishes from Bergeron et al. (2023) (Fig. 3e), we found no significant correlation between effective population size and mutation rate. This lack of correlation was robust after phylogenetic correction using the fish tree of life from Rabosky et al. (2018) for these six fish species, which all come from different families. Furthermore, three out of the six fish mutation rates from Bergeron et al. (2023) were higher than our Devils Hole pupfish estimate despite their much larger effective population sizes (Fig. 3e).

### Mutation spectra

The overall mutational spectrum of *de novo* mutations within *C. diabolis* was dominated by C>T mutations (38.3%) with 46.2% at CpG sites (Fig. 4b). This spectrum is very similar to the averaged spectra across 8 species of fish from Bergeron et al. (2023), in which the predominant type was also C>T mutations (46.7%) with 44.5% at CpG sites (Fig. 4a).

**Fig. 4.**
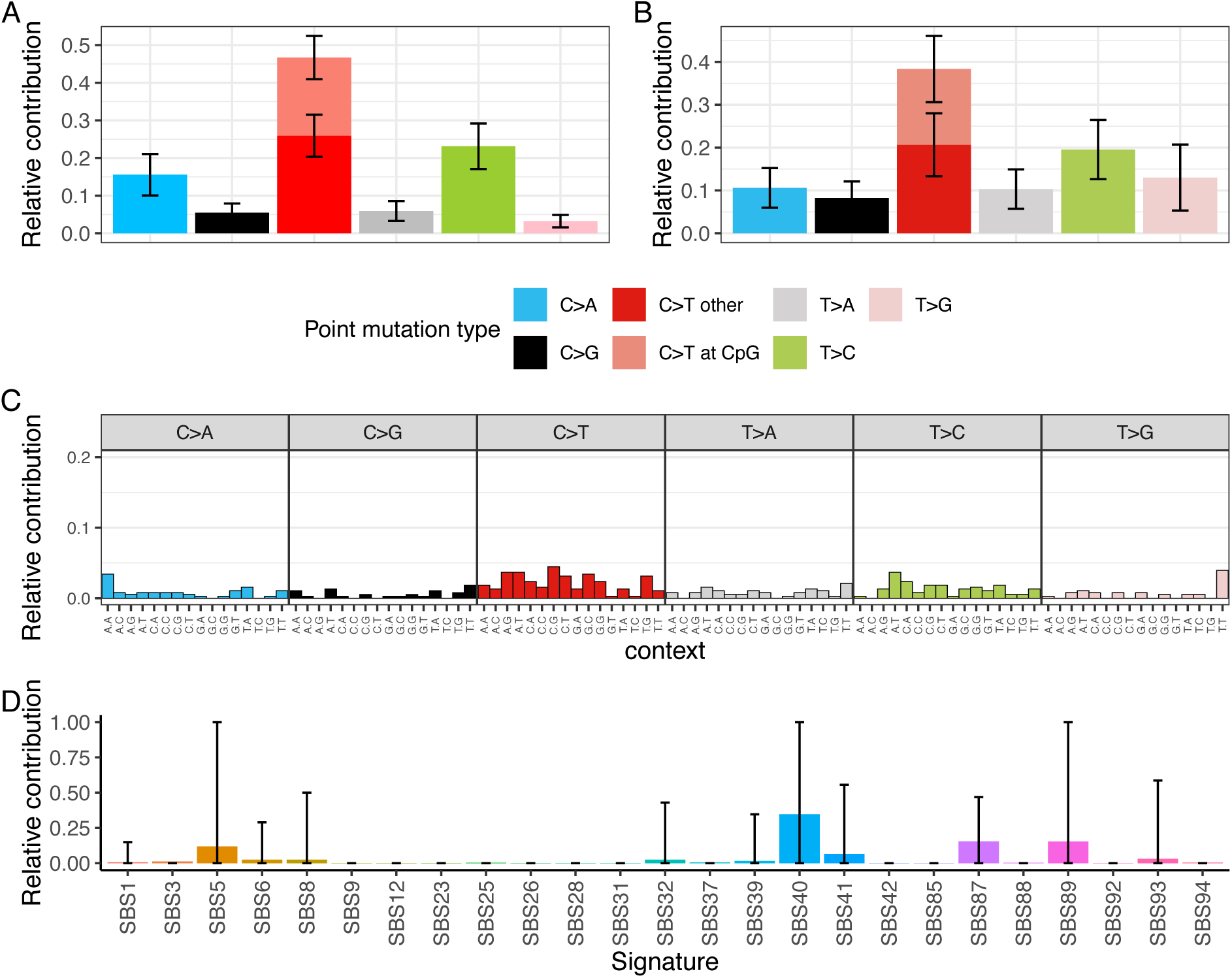
Mutational spectra and signatures. (a) Single base mutation spectra averaged across 89 mutations from eight fish species from Bergeron et al. (2023) and (b) for *C. diabolis*. Error bars represent 95% confidence interval. (c) 96 tri-nucleotide mutation spectra of *C. diabolis.* (d) *C. diabolis* mutational signature fitting using MutationalPatterns (Blokzijl et al. 2018). Signatures were refitted with max delta = 0.01 to avoid overfitting and 1,000 bootstraps to assess the signature refitting stability. Error bars represent 95% confidence interval.

We compared single base pair mutation spectra between mature adults and embryonic lethals and found them to be similar (Supp. Fig. 1). We also analyzed the 96-trinucleotide mutation spectrum and found a significant difference in the relative contribution of ATT ◊ AGT mutations between mature adults (4 mutations) and embryonic lethals (0 mutations) (*P* = 0.004, Fisher’s exact test) (Supp. Fig. 2; although this result was not significant following multiple test correction: Benjamini-Hochberg adjusted *P* = 0.318). The following mutational signatures were inferred to make the largest contributions to the overall trinucleotide spectrum: SBS40, associated with gamma irradiation (Manaka et al. 2024) and DNA repair deficiencies that trigger translesion synthesis (Hwang et al. 2025); SBS41, unknown aetiology (Alexandrov et al. 2020); SBS87, associated with thiopurine chemotherapy (Li et al. 2020); and SBS89, unknown aetiology (Lee-Six et al. 2019) (Fig. 4d, Supp. Fig. 3).

## Discussion

We estimated a mutation rate of 8.09 × 10^−9^ mutations per base pair per generation within the Devils Hole pupfish, taking advantage of their high levels of inbreeding to measure recent mutations within autozygous segments for 58 medium-coverage (mean = 18X) and four high-coverage (mean = 98X) individuals. Devils Hole pupfish has a mutation rate that is lower than predicted by the drift-barrier hypothesis based on their extremely low effective population size (Fig. 3d) (Lynch 2010; Sung et al. 2012; Lynch et al. 2016; Lynch et al. 2023). When restricted to actinopterygian fishes, we found no relationship between effective population size and mutation rate. Furthermore, 3 out of 6 fish species with much higher population sizes had higher estimated mutation rates than the Devils Hole pupfish, in direct contrast to predictions of the drift-barrier hypothesis. Overall, our results call into question the drift-barrier hypothesis as a robust explanation for variation in mutation rates across taxa and are consistent with recent results that suggest no relationship between effective population size and mutation rate when accounting for differences in generation times across taxa (Weinstein and Roy 2026).

### Tentative support for the drift-barrier hypothesis

While our empirical comparisons to other vertebrates are limited by lack of data at the lower extreme, the directional patterns appear consistent with the drift-barrier model and Devils Hole pupfish have a higher mutation rate than the mean rate in other fishes (Bergeron et al. 2023). A recent meta-analysis of fish mutation rates by Zhang et al. (2025) found only three fish species with higher mutation rates than Devils Hole pupfish. However, the mutation rate estimates for Japanese flounder (8.54 × 10^−9^ per bp per generation) and tongue sole (9.13 × 10^−9^ per bp per generation) were based on sequencing of only two or three trios which introduces substantial uncertainty, and were derived from domesticated sources, which often have elevated mutation rates (Bergeron et al. 2023). Furthermore, the mean mutation rate estimate for the third species (guppy: 9.28 × 10^−9^ per bp per generation) included data from one family that had a 7-fold higher outlier mutation rate (2.90 × 10^−8^ per bp per generation) relative to mean rates across four other guppy families (4.17 × 10^−9^ per bp per generation) (Burda and Konczal 2023; Lin et al. 2023; Zhang et al. 2025). A recent meta-analysis of animal germline mutation rates by de Manuel and Prezworski (2025) that only included mutation rate estimates of species with at least five trios suggests that mutation rate estimates among fishes are quite low, ranging from 2.00 × 10^−9^ – 5.05 × 10^−9^ per bp per generation, and they reported a guppy mutation rate estimate of only 3.4 × 10^−9^ per bp per generation. Lower mutation rates among teleost fishes may suggest specific genetic or physiological mechanisms regulating mutation rates in this group, although there are currently a limited number of fish mutation rate estimates in the literature.

Another possibility is that small effective population size has increased genomic stress and burden within Devils Hole pupfish in ways consistent with the drift-barrier hypothesis, but this is not yet reflected in the base substitution mutation rate. Previous work hypothesized an elevated mutation rate in Devils Hole pupfish in part due to physiological stresses (Martin and Höhna 2018), and here we find a contribution of the SBS40 mutational signature to the Devils Hole pupfish mutation spectrum, which is associated with DNA repair deficiencies and genomic instability (Hwang et al. 2025). However, several studies have shown that physiological stress tends to increase the indel mutation rate with little to no effect on the base substitution mutation rate, such as in green alga subjected to salinity stress (Hasan et al. 2022), *A. thaliana* under thermal stress (Belfield et al. 2021), and *C. elegans* exposed to increased cellular oxidative stress (Rajaei et al. 2021). In addition, a recent study found a negative correlation between effective population size and alternative splicing rates, and that drift can reduce the capacity of selection to prevent splicing errors (Bénitière et al. 2024). Finally, it is unclear on what timescales mutation rates evolve and whether the current mutation rate has rapidly evolved since initial colonization of Devils Hole. Potentially Devils Hole pupfish are too young for the mutation rate to have drifted substantially higher.

### Adult vs. embryonic lethal mutation rate

The elevated mutation rate among embryonic lethal individuals compared to adults is consistent with the idea that mutation rates are sensitive to genetic stress (Sharp and Agrawal 2012). The difference in mutation rate may suggest the presence of segregating lethal mutator alleles in the population, consistent with the expectation that species with short generation times and low effective population sizes have reduced strength of selection against mutator alleles (Zhu et al. 2025). Mutator alleles in embryonic lethal samples could increase the mutation rate past the point of viability, thus preventing the detection of elevated mutation rates in adults.

However, while adults were sequenced from one tissue (fin clips), entire embryos were sequenced for our embryonic lethal samples, and mutation rates can vary substantially among somatic cell types due to differing selective pressures (de Manuel et al. 2025). Thus, our embryonic lethal estimate may be inflated by the accidental inclusion of additional somatic mutations from multiple tissue types. Finally, the difference in mutation rate between adult and embryonic lethal individuals may be a byproduct of dying embryos rapidly accumulating more somatic mutations after the loss of cellular homeostasis.

### Mutational spectra

We found that Devils Hole pupfish mutation spectra were relatively similar to other fishes. When we analyzed the 96-trinucleotide mutation spectrum, we found no significant differences between our embryonic lethal and adult samples, despite the higher mutation rates observed in embryonic lethal individuals. This increase in embryos that died prematurely may reflect an endogenously-driven increase in overall mutation rate rather than an environmentally-driven increase that would leave specific patterns in the mutational spectra. Alternatively, our low sample size of mutations may severely limit our ability to infer mutational signatures in the spectrum.

We had limited power to infer mutational signatures. Nevertheless, the following signatures were observed to make the largest contributions to the overall spectra profile: SBS40, associated with gamma irraditaion (Manaka et al. 2024) and DNA repair deficiencies that trigger translesion synthesis (Hwang et al. 2025), SBS41, unknown aetiology (Alexandrov et al. 2020), SBS87, associated with thiopurine chemotherapy (Li et al. 2020), and SBS89, unknown aetiology (Lee-Six et al. 2019) (Fig. 4d, Supp. Fig. 3). The SBS40 signature reflects potential defects in DNA repair machinery. Although we had limited power to fit signatures to our mutational spectra, we observed that SBS40 may contribute to our spectra, consistent with the drift-barrier expectation that increased mutation rates result from mutator alleles drifting to higher frequency in the population (Lynch 2008).

Intriguingly, there is a segregating partial deletion of RAD23A in Devils Hole pupfish (Tian et al. 2026). RAD23A is a nucleotide excision repair protein that interacts with DNA polymerases in the translesion synthesis pathway (Ashton et al. 2023); future investigation is needed to functionally test if this deletion is contributing to this elevated SBS40 signature and resulting in higher mutation rates in Devils Hole pupfish.

### Caveats: Potential errors in mutation rate analysis

It is well known that mutation rate estimates can vary up to 2-fold even due to different bioinformatics pipelines and that these estimates are highly sensitive to coverage and numerous biases such as genomic sequencing library prep methods and sample tissue type (Bergeron et al. 2022). In addition, as the quality of genomic data has increased, many mutation rates have decreased significantly, for example in cichlids (6.6 × 10^−8^ per bp per generation (Recknagel et al. 2013) vs. 3.5 × 10^−9^ per bp per generation (Malinsky et al. 2018)). Genomic estimates of *N_e_* based on our estimated mutation rate (*N_e_* = 426) are similar but slightly above our estimates of *N_e_* based on the harmonic mean of wild census population size over time (spring = 122, fall = 209) (Tian et al. 2022), suggesting that our mutation rate is within a reasonable range but could be an underestimate. Previous studies have shown that singletons have a higher false negative rate than SNPs estimated at higher allele frequencies (above 10%), due to the nature of multi-sample variant calling methods using the entire dataset to make calls (Narasimhan et al. 2017), which could also result in underestimation of the mutation rate.

Overestimation of the Devils Hole pupfish mutation rate is possible if not all detected singletons were truly from the previous generation, which would bias our estimated mutation rate upwards. Our singleton-based estimates were higher than our gene-conversion corrected estimates. The singleton-based estimate could be biased upwards due to accidental inclusion of gene conversions from rare alleles. Meanwhile, the gene conversion corrected estimate may be lower due to underestimation of the number of meioses or the removal of recurrent mutations.

The confidence interval for the drift-barrier prediction was quite large due to extensive variation among taxa and a lack of data at the extremes. Although Bergeron et al. (2023) reported 40-fold variation in vertebrate mutation rates, a recent review that conservatively only incorporated mutation rates for species with a minimum of five trios suggests that vertebrate mutation rates may only vary by one order of magnitude (de Manuel et al. 2025).

### Conservation prospects for the Devils Hole pupfish

Our estimates of effective population size in Devils Hole pupfish are slightly higher than the recommended minimum to limit inbreeding depression (*N_e_* > 100) (Frankham et al. 2014), although there is evidence of inbreeding depression in the population (Tian et al. 2026). In addition, *N_e_* is far below the recommended minimums for maintaining evolutionary potential (*N_e_* > 1000) (Frankham et al. 2014), suggesting that despite recovery of the population following bottlenecks in 2007 and 2013 and recent growth from an all-time low of 20 individuals in 2025, the species remains at risk and will benefit from continued conservation and recovery efforts.

Our study highlights the possibilities for estimating mutation rates in endangered species and illustrates how this can be done even when pedigree-based methods are unavailable or impossible. This will facilitate further testing of the drift-barrier hypothesis at an expanded range of *N_e_* values and improve our understanding of the factors that drive variation in mutation rates across the tree of life.

## Methods

### Sampling

We analyzed sequenced whole genomes available on NCBI (mean coverage 18X, range = 15X-25X) (Tian et al. 2026) aligned to a chromosome-level *de novo* genome assembly for *C. diabolis* (Genbank: GCA_030533445.1) (Tian et al. 2026). We chose 58 samples with at least 15X coverage, given that lower coverage strongly biases autozygous segment lengths, leading to higher rates of false positives and negatives. Sequencing of adults was based on caudal fin clips collected from adult fish reared in the refuge facility (Ash Meadows Fish Conservation Facility) from wild-collected or refuge-collected eggs and stored in 100% EtOH at −20 °C. Sequencing of embryonic lethal individuals used DNA extracted only from living embryos visibly showing phenotypic signs of developmental slowdown, such as a heart defect at 5-7 days post fertilization, and stored in 100% EtoH at −20 °C. For an additional 4 samples (RT10, RT11, RT14, RT17) that were selected due to their high levels of relatedness inferred using KING (Manichaikul et al. 2010), we performed additional sequencing for this study (mean coverage 98X, range = 88-113X), resulting in a total of 62 individuals.

### Whole genome sequencing and variant calling

Raw reads were trimmed with fastp (v0.23.4) (Chen et al. 2018). Reads were mapped to the UCB_CyDiab_1.0 *Cyprindon diabolis* reference genome (GCA_030533445.1) with bwa mem (v0.7.17) (Li 2013). Duplicate reads in the bam files were marked with Picard MarkDuplicates (GATK v4.6.1.0) (Broad Institute). Coverage and mapping quality were assessed with qualimap (v2.2.2) (Okonechnikov et al. 2016). We followed the GATK (v.4.6.1.0) (McKenna et al. 2010) genotyping pipeline to call variants. We used HaplotypeCaller to call variants and stored variants in the GenomicsDB datastore format prior to using GenotypeGVCFs to perform joint genotyping. We restricted our analyses to biallelic SNPs and applied the recommended GATK hard filters (QD < 2, QUAL < 30, SOR > 3, FS > 60, MQ < 40, MQRankSum < −12.5, ReadPosRankSum < −8) (DePristo et al. 2011; Van der Auwera et al. 2013) to conservatively call SNPs and then used this set of SNPs to perform BQSR. After recalibrating the bam files, we followed subsequent steps and went through another round of variant calling, joint genotyping, and hard filtering. These additional filters were applied: We ensured that SNPs had to be in regions of the genome where reads mapped uniquely, based on a mappability score of greater than 0.5 from h (-K 150, -E 2) (v1.3.0) (Pockrandt et al. 2020), a genotype quality of at least 30, a maximum missingness of 25%, and depth not exceeding the 99^th^ percentile of total depth across individuals using bcftools (v1.16) (Danecek et al. 2021) and vcftools (v0.1.16) (Danecek et al. 2011). At an individual level, we also removed sites with less than 1/3X coverage or greater than 2X coverage based on individual coverage. This resulted in a final set of 343,487 high-quality SNPs, that we used to call autozygous segments.

To identify putative mutations, we applied even more stringent additional filters to increase confidence in the detection of true de novo mutations, followed the recommendations of Wu et al. (2020). We required mutations to have a genotype quality of at least 40 and a quality score of at least 100. We also required the number of reads supporting the alternate mutant allele to be at least 4. Finally, for each SNP, we performed a 2-sided binomial test on allelic balance (number of reads supporting alternate allele / total reads) and filtered SNPs with P-value below 0.05. With a frequency of 0.5, we assumed that mutations were germline and not mosaic or somatic.

### Identification of autozygous segments

Autozygous segments were identified using BCFtools/RoH with the Genotypes method (Narasimhan et al. 2016). BCFtools/RoH is more conservative than other methods and tends to have lower false positive rates but higher false negative rates, which is more appropriate for studies such as this one that require higher accuracy (Silva et al. 2024). We used genotypes with a genotype quality of at least 30 to infer 2,754 autozygous segments. We restricted our analyses to segments at least 5 Mb long, to focus on segments that originated in a very recent common ancestor, leaving 526 segments, totaling 2.99 Gb, across 42 individuals. For each segment, we excluded 1 Mb from each end to avoid calling mutations in regions adjacent to autozygous segments with a higher TMRCA, following the procedure in Narasimhan et al. (2017).

### Estimation of de novo mutation rates

We calculated mutation rate as the number of heterozygous SNPs found within autozygous segments with 1 Mb ends cut off (N) divided by the length of callable basepairs within these portions of the autozygous segments (L) and the number of meioses (M): N / (L*M), following Narasimhan et al. (2017). The number of callable basepairs was calculated within the analyzed autozygous segments with the same mappability mask used for SNP filtering.

To account for gene conversions, we applied a minor allele frequency threshold approach to calculate the mutation rate with varying levels of maximum allowed MAF for heterozygotes in our autozygous segments, and inferred the y-intercept as the gene-conversion corrected mutation rate (Palamara et al. 2015). Following Palamara et al. (2015), we estimated the gene conversion rate as the difference between the uncorrected and corrected mutation rates, divided by the population heterozygosity (*H* = 1.38 × 10^−5^) (Tian et al. 2026).

Lacking a pedigree, we were unable to directly calculate the number of meioses separating autozygous segments from their TMRCA. Consequently, we estimated the number of meioses based on the expectation that a separation of M meioses generates segment lengths that are exponentially distributed with a mean of 100/M cM (Thompson 2013). To convert our segment lengths in Mb to cM, we used both a genome-wide average recombination rate of ∼2.8 cM/Mb (1.4 – 6.3 cM/Mb, 25^th^ – 75^th^ percentiles) and the median genome-wide recombination rate for fishes (Stapley et al. 2017). This resulted in a weighted mean of 7.54 meioses across our dataset (7.19 for embryonic lethals, 8.35 for mature adults). We weighted the estimated number of meioses separating chromosomes in individuals by the proportion of the genome with autozygous segments larger than 5 Mb in each individual. Across individuals, the estimated number of meioses ranged from 6.2 – 15.9 (Fig. 1).

We also calculated mutation rate estimates based on singletons only, which are expected to have arisen in the preceding generation. This mutation rate was calculated as N = number of singletons, L = callable basepairs, and M = 2. For embryonic lethal and mature adult groups, we bootstrapped resampled individuals with replacement (10,000 replicates). For each bootstrap replicate, we estimated the singleton mutation rate as the total number of singletons across sampled individuals divided by twice the total callable bp to generate 95% confidence intervals for the mean singleton mutation rates and difference between groups.

### Comparison to vertebrate species

We compared *C. diabolis* mutation rates to other vertebrate species in Bergeron et al. (2023) for generation time, age at sexual maturity, lifespan in the wild, and effective population size, all factors correlated with *de novo* mutation rates. For *C. diabolis*, we used a generation time of 9 months, an age at sexual maturity of four months, and a lifespan in the wild of one year (Deacon et al. 1995). For effective population size, we used an *N_e_* of 165, which is the mean of the spring and fall effective population sizes calculated from the harmonic mean of census population size from 1970 – 2020. This approach differs from Bergeron et al. (2023) which relied on PSMC population size estimates from 30,000 – 1,000,000 years ago because *C. diabolis* is far younger. However, the harmonic mean of population size estimated from PSMC is significantly correlated with effective population size estimated from the equation *N_e_* = π/4µ (Bergeron et al. 2023). This approach yields slightly higher estimates of *N_e_* (426), which we also used to predict a mutation rate based on the drift-barrier hypothesis.

### Mutational spectra

We analyzed the 6-base substitution and 96-trinucleotide mutational spectra of our 380 identified mutations using MutationalPatterns (Blokzijl et al. 2018; Manders et al. 2022) and compared them to the mutational spectra of 8 fishes from Bergeron et al. (2023). We used the “fit_to_signatures” function in MutationalPatterns to identify the optimal combination of mutational signatures that best reconstructs our observed mutation spectra. This often results in many signatures being used to explain mutational spectra due to overfitting. As a result, we used the “fit_to_signatures_strict” function to refit signatures and sequentially remove signatures with the lowest contribution to the overall profile until the cosine similarity between the original and refitted profile is greater than the “max_delta” parameter, which we set to 0.01. It is difficult to infer whether differences between embryonic lethals and adults in relative proportions of signatures are meaningful or potentially due to our small sample sizes and model overfitting. To assess how robust our refitted mutation profiles were, we used the “fit_to_signatures_bootstrapped” function to bootstrap sample mutations with replacement using the original mutation profile as weights 1,000 times, performing a signature refitting at each iteration, with max_delta set to 0.01.

## Data availability

All novel sequence data generated by this study is available at NCBI’s SRA database under XXX. Genome assemblies are available on NCBI. Code for all analyses is available on the first author’s github account at https://github.com/tiandavid/mutation_rate.

## Acknowledgements

This study was funded by the USDI Fish and Wildlife Service F20AC10915-01, the Hellman Foundation faculty award, and NSF CAREER 1749764 to C.H.M. D.T. was supported by an NSF GRFP (DGE 1752814) and a Philomathia Graduate Fellowship in the Environmental Sciences. We thank the National Park Service and US Fish & Wildlife for their work in conserving this species. The findings and conclusions in this article are those of the authors and do not necessarily represent the views of the U.S. Fish and Wildlife Service or National Park Service. We thank members of the Martin lab for valuable feedback and discussion of this project.

## Author Contributions

D.T., P.M. and C.H.M. conceptualized the project. D.T. performed all analyses. E.K. assisted with methodology. C.H.M. provided samples and funding for sequencing and analyses. P.M. was supported by the Burroughs Wellcome Fund (Career Award at the Scientific Interface). D.T. wrote the manuscript. K.W., O.F., M.S., and J.G. collected samples. All authors edited and reviewed the manuscript.

## References

Alexandrov LB et al. 2020. The repertoire of mutational signatures in human cancer. Nature. 578(7793):94–101. 10.1038/s41586-020-1943-3

Ashton NW et al. 2023. A Novel Interaction Between RAD23A/B and Y-family DNA Polymerases. Journal of Molecular Biology. 435(24):168353. 10.1016/j.jmb.2023.168353

Baer CF, Miyamoto MM, Denver DR. 2007. Mutation rate variation in multicellular eukaryotes: causes and consequences. Nat Rev Genet. 8(8):619–631. 10.1038/nrg2158

Barton NH, Keightley PD. 2002. Understanding quantitative genetic variation. Nat Rev Genet. 3(1):11–21. 10.1038/nrg700

Belfield EJ et al. 2021. Thermal stress accelerates Arabidopsis thaliana mutation rate. Genome Res. 31(1):40–50. 10.1101/gr.259853.119

Bénitière F, Necsulea A, Duret L. 2024. Random genetic drift sets an upper limit on mRNA splicing accuracy in metazoans Castric V, Perry GH, editors. eLife. 13:RP93629. 10.7554/eLife.93629

Bergeron LA et al. 2022. The Mutationathon highlights the importance of reaching standardization in estimates of pedigree-based germline mutation rates Courtier-Orgogozo V, Perry GH, Quinlan AR, editors. eLife. 11:e73577. 10.7554/eLife.73577

Bergeron LA, et al. 2023. Evolution of the germline mutation rate across vertebrates. Nature. 615(7951):285–291. 10.1038/s41586-023-05752-y

Blokzijl F, Janssen R, van Boxtel R, Cuppen E. 2018. MutationalPatterns: comprehensive genome-wide analysis of mutational processes. Genome Medicine. 10(1):33. 10.1186/s13073-018-0539-0

Broad Institute. [accessed 2021 May 6]. http://broadinstitute.github.io/picard/

Burda K, Konczal M. 2023. Validation of machine learning approach for direct mutation rate estimation. Molecular Ecology Resources. 23(8):1757–1771. 10.1111/1755-0998.13841

Cadet J, Wagner JR. 2013. DNA Base Damage by Reactive Oxygen Species, Oxidizing Agents, and UV Radiation. Cold Spring Harb Perspect Biol. 5(2):a012559. 10.1101/cshperspect.a012559

Campbell CD et al. 2012. Estimating the human mutation rate using autozygosity in a founder population. Nat Genet. 44(11):1277–1281. 10.1038/ng.2418

Chao L, Carr DE. 1993. The Molecular Clock and the Relationship between Population Size and Generation Time. Evolution. 47(2):688–690. 10.2307/2410082

Chen S, Zhou Y, Chen Y, Gu J. 2018. fastp: an ultra-fast all-in-one FASTQ preprocessor. Bioinformatics. 34(17):i884–i890. 10.1093/bioinformatics/bty560

Chintalapati M, Moorjani P. 2020. Evolution of the mutation rate across primates. Current Opinion in Genetics & Development. 62:58–64 (Genetics of Human Origin). 10.1016/j.gde.2020.05.028

Danecek P et al. 2011. The variant call format and VCFtools. Bioinformatics. 27(15):2156–2158. 10.1093/bioinformatics/btr330

Danecek P et al. 2021. Twelve years of SAMtools and BCFtools. GigaScience. 10(2):giab008. 10.1093/gigascience/giab008

Deacon JE. 1979. Endangered and Threatened Fishes of the West. Great Basin Naturalist Memoirs. (3):41–64

Deacon JE, Taylor FR, Pedretti JW. 1995. Egg viability and ecology of Devils Hole pupfish: Insights from captive propagation. The Southwestern Naturalist. 40(2):216–223

DePristo MA et al. 2011. A framework for variation discovery and genotyping using next-generation DNA sequencing data. Nat Genet. 43(5):491–498. 10.1038/ng.806

Feng C et al. 2017. Moderate nucleotide diversity in the Atlantic herring is associated with a low mutation rate Przeworski M, editor. eLife. 6:e23907. 10.7554/eLife.23907

Foote AD et al. 2021. Runs of homozygosity in killer whale genomes provide a global record of demographic histories. Molecular Ecology. 30(23):6162–6177. 10.1111/mec.16137

Frankham R, Bradshaw CJA, Brook BW. 2014. Genetics in conservation management: Revised recommendations for the 50/500 rules, Red List criteria and population viability analyses. Biological Conservation. 170:56–63. 10.1016/j.biocon.2013.12.036

Gao Z et al. 2019. Overlooked roles of DNA damage and maternal age in generating human germline mutations. Proceedings of the National Academy of Sciences. 116(19):9491–9500. 10.1073/pnas.1901259116

Gustafson ES, Deacon JE. 1997. Distribution of larval Devils Hole pupfish, Cyprinodon diabolis Wales, in relation to dissolved oxygen concentration in Devil’s Hole. Report to the National Park Service, Death Valley National Park.

Hasan AR et al. 2022. Salt stress alters the spectrum of de novo mutation available to selection during experimental adaptation of Chlamydomonas reinhardtii. Evolution. 76(10):2450–2463. 10.1111/evo.14604

Hoeijmakers JHJ. 2009. DNA Damage, Aging, and Cancer. New England Journal of Medicine. 361(15):1475–1485. 10.1056/NEJMra0804615

Hwang T et al. 2025. Comprehensive whole-genome sequencing reveals origins of mutational signatures associated with aging, mismatch repair deficiency and temozolomide chemotherapy. Nucleic Acids Res. 53(1):gkae1122. 10.1093/nar/gkae1122

James CJ. 1969. Aspects of the Ecology of the Devil’s Hole Pupfish (Cyprinodon Diabolis) Wales. University of Nevada, Las Vegas.

Jeffreys AJ, Royle NJ, Wilson V, Wong Z. 1988. Spontaneous mutation rates to new length alleles at tandem-repetitive hypervariable loci in human DNA. Nature. 332(6161):278–281. 10.1038/332278a0

Kessler MD et al. 2020. De novo mutations across 1,465 diverse genomes reveal mutational insights and reductions in the Amish founder population. Proceedings of the National Academy of Sciences. 117(5):2560–2569. 10.1073/pnas.1902766117

Kimura M. 1967. On the evolutionary adjustment of spontaneous mutation rates. Genetics Research. 9(1):23–34. 10.1017/S0016672300010284

Lee-Six H et al. 2019. The landscape of somatic mutation in normal colorectal epithelial cells. Nature. 574(7779):532–537. 10.1038/s41586-019-1672-7

Li B et al. 2020. Therapy-induced mutations drive the genomic landscape of relapsed acute lymphoblastic leukemia. Blood. 135(1):41–55. 10.1182/blood.2019002220

Li H. 2013. Aligning sequence reads, clone sequences and assembly contigs with BWA-MEM. http://arxiv.org/abs/1303.3997. 10.48550/arXiv.1303.3997

Lin Y et al. 2023. Extensive variation in germline de novo mutations in Poecilia reticulata. Genome Res. 33(8):1317–1324. 10.1101/gr.277936.123

Liu H, Zhang J. 2019. Yeast Spontaneous Mutation Rate and Spectrum Vary with Environment. Current Biology. 29(10):1584–1591.e3. 10.1016/j.cub.2019.03.054

Lynch M. 2008. The Cellular, Developmental and Population-Genetic Determinants of Mutation-Rate Evolution. Genetics. 180(2):933–943. 10.1534/genetics.108.090456

Lynch M. 2010. Evolution of the mutation rate. Trends in Genetics. 26(8):345–352. 10.1016/j.tig.2010.05.003

Lynch M et al. 2016. Genetic drift, selection and the evolution of the mutation rate. Nat Rev Genet. 17(11):704–714. 10.1038/nrg.2016.104

Lynch M et al. 2023. The divergence of mutation rates and spectra across the Tree of Life. EMBO reports. 24(10):e57561. 10.15252/embr.202357561

Malinsky M et al. 2018. Whole-genome sequences of Malawi cichlids reveal multiple radiations interconnected by gene flow. Nat Ecol Evol. 2(12):1940–1955. 10.1038/s41559-018-0717-x

Manaka Y et al. 2024. Single base substitution signatures 17a, 17b, and 40 are induced by γ-ray irradiation in association with increased reactive oxidative species. Heliyon. 10(6):e28044. 10.1016/j.heliyon.2024.e28044

Manders F et al. 2022. MutationalPatterns: the one stop shop for the analysis of mutational processes. BMC Genomics. 23(1):134. 10.1186/s12864-022-08357-3

Manichaikul A et al. 2010. Robust relationship inference in genome-wide association studies. Bioinformatics. 26(22):2867–2873. 10.1093/bioinformatics/btq559

de Manuel M, Przeworski M, Spisak N, Anastasia Stolyarova. 2025. What sets the mutation rate of a cell type in an animal species? 2025.12.19.695482 [accessed 2026 Jan 8]. https://www.biorxiv.org/content/10.64898/2025.12.19.695482v1. 10.64898/2025.12.19.695482

Martin AP, Palumbi SR. 1993. Body size, metabolic rate, generation time, and the molecular clock. Proceedings of the National Academy of Sciences. 90(9):4087–4091. 10.1073/pnas.90.9.4087

Martin CH et al. 2017. The complex effects of demographic history on the estimation of substitution rate: concatenated gene analysis results in no more than twofold overestimation. Proceedings of the Royal Society B: Biological Sciences. 284(1860):20170537. 10.1098/rspb.2017.0537

Martin CH, Crawford Jacob E., Turner Bruce J., Simons Lee H. 2016. Diabolical survival in Death Valley: recent pupfish colonization, gene flow and genetic assimilation in the smallest species range on earth. Proceedings of the Royal Society B: Biological Sciences. 283(1823):20152334. 10.1098/rspb.2015.2334

Martin CH, Höhna S. 2018. New evidence for the recent divergence of Devil’s Hole pupfish and the plausibility of elevated mutation rates in endangered taxa. Mol Ecol. 27(4):831–838. 10.1111/mec.14404

McKenna A et al. 2010. The Genome Analysis Toolkit: A MapReduce framework for analyzing next-generation DNA sequencing data. Genome Research. 20(9):1297. 10.1101/gr.107524.110

Miller RR. 1948. The Cyprinodont fishes of the Death Valley system of eastern California and southwestern Nevada. undefined. [published online ahead of print] [accessed 2021 June 30]

Mugal CF, Arndt PF, Holm L, Ellegren H. 2015. Evolutionary Consequences of DNA Methylation on the GC Content in Vertebrate Genomes. G3 Genes|Genomes|Genetics. 5(3):441–447. 10.1534/g3.114.015545

Narasimhan V et al. 2016. BCFtools/RoH: a hidden Markov model approach for detecting autozygosity from next-generation sequencing data. Bioinformatics. 32(11):1749–1751. 10.1093/bioinformatics/btw044

Narasimhan VM et al. 2017. Estimating the human mutation rate from autozygous segments reveals population differences in human mutational processes. Nat Commun. 8(1):303. 10.1038/s41467-017-00323-y

Ohta T. 1993. An examination of the generation-time effect on molecular evolution. Proceedings of the National Academy of Sciences. 90(22):10676–10680. 10.1073/pnas.90.22.10676

Okonechnikov K, Conesa A, García-Alcalde F. 2016. Qualimap 2: advanced multi-sample quality control for high-throughput sequencing data. Bioinformatics. 32(2):292–294. 10.1093/bioinformatics/btv566

Palamara PF et al. 2015. Leveraging Distant Relatedness to Quantify Human Mutation and Gene-Conversion Rates. The American Journal of Human Genetics. 97(6):775–789. 10.1016/j.ajhg.2015.10.006

Peña-García Y et al. 2025. Low mutation rate but high male-bias in the germline of a short-lived opossum. Genetics. iyaf177. 10.1093/genetics/iyaf177

Pockrandt C, Alzamel M, Iliopoulos CS, Reinert K. 2020. GenMap: ultra-fast computation of genome mappability. Bioinformatics. 36(12):3687–3692. 10.1093/bioinformatics/btaa222

Poetsch AR, Boulton SJ, Luscombe NM. 2018. Genomic landscape of oxidative DNA damage and repair reveals regioselective protection from mutagenesis. Genome Biology. 19(1):215. 10.1186/s13059-018-1582-2

Rabosky DL et al. 2018. An inverse latitudinal gradient in speciation rate for marine fishes. Nature. 559(7714):392–395. 10.1038/s41586-018-0273-1

Rajaei M et al. 2021. Mutability of mononucleotide repeats, not oxidative stress, explains the discrepancy between laboratory-accumulated mutations and the natural allele-frequency spectrum in C. elegans. Genome Res. 31(9):1602–1613. 10.1101/gr.275372.121

Recknagel H, Elmer KR, Meyer A. 2013. A Hybrid Genetic Linkage Map of Two Ecologically and Morphologically Divergent Midas Cichlid Fishes (Amphilophus spp.) Obtained by Massively Parallel DNA Sequencing (ddRADSeq). G3 Genes|Genomes|Genetics. 3(1):65–74. 10.1534/g3.112.003897

Riggs AC, Deacon JE. 2002. Connectivity in Desert Aquatic Ecosystems: The Devils Hole Story. 38

Sharp NP, Agrawal AF. 2012. Evidence for elevated mutation rates in low-quality genotypes. Proceedings of the National Academy of Sciences. 109(16):6142–6146. 10.1073/pnas.1118918109

Silva GAA et al. 2024. Detectability of runs of homozygosity is influenced by analysis parameters and population-specific demographic history. PLOS Computational Biology. 20(10):e1012566. 10.1371/journal.pcbi.1012566

Stamatoyannopoulos JA et al. 2009. Human mutation rate associated with DNA replication timing. Nat Genet. 41(4):393–395. 10.1038/ng.363

Stapley J et al. 2017. Recombination: the good, the bad and the variable. Philosophical Transactions of the Royal Society B: Biological Sciences. 372(1736):20170279. 10.1098/rstb.2017.0279

Stoike SL, Pister EP. 2010. Threatened fishes of the world: Cyprinodon diabolis (Wales, 1930). Environ Biol Fish. 88(4):399–400. 10.1007/s10641-010-9651-8

Sung W et al. 2012. Drift-barrier hypothesis and mutation-rate evolution. Proceedings of the National Academy of Sciences. 109(45):18488–18492. 10.1073/pnas.1216223109

Thomas GWC et al. 2018. Reproductive Longevity Predicts Mutation Rates in Primates. Current Biology. 28(19):3193–3197.e5. 10.1016/j.cub.2018.08.050

Thompson EA. 2013. Identity by Descent: Variation in Meiosis, Across Genomes, and in Populations. Genetics. 194(2):301–326. 10.1534/genetics.112.148825

Tian D et al. 2026. Long-term small effective population size, inbreeding, and a recessive lethal haplotype drive premature death in the endangered Devils Hole pupfish (Cyprinodon diabolis). 2026.06.06.730634 [accessed 2026 June 23]. https://www.biorxiv.org/content/10.64898/2026.06.06.730634v2. 10.64898/2026.06.06.730634

Tian D, Patton AH, Turner BJ, Martin CH. 2022. Severe inbreeding, increased mutation load and gene loss-of-function in the critically endangered Devils Hole pupfish. Proceedings of the Royal Society B: Biological Sciences. 289(1986):20221561. 10.1098/rspb.2022.1561

Van der Auwera GA et al. 2013. From FastQ Data to High-Confidence Variant Calls: The Genome Analysis Toolkit Best Practices Pipeline. Current Protocols in Bioinformatics. 43(1):11.10.1–11.10.33. 10.1002/0471250953.bi1110s43

Volkova NV et al. 2020. Mutational signatures are jointly shaped by DNA damage and repair. Nat Commun. 11(1):2169. 10.1038/s41467-020-15912-7

Wang RJ et al. 2022. Examining the Effects of Hibernation on Germline Mutation Rates in Grizzly Bears. Genome Biol Evol. 14(10):evac148. 10.1093/gbe/evac148

Wang Y, Obbard DJ. 2023. Experimental estimates of germline mutation rate in eukaryotes: a phylogenetic meta-analysis. Evolution Letters. 7(4):216–226. 10.1093/evlett/qrad027

Waples RS, Luikart G, Faulkner JR, Tallmon DA. 2013. Simple life-history traits explain key effective population size ratios across diverse taxa. Proceedings of the Royal Society B: Biological Sciences. 280(1768):20131339. 10.1098/rspb.2013.1339

Wayne RK et al. 1991. A MORPHOLOGIC AND GENETIC STUDY OF THE ISLAND FOX, UROCYON LITTORALIS. Evol. 45(8):1849–1868. 10.1111/j.1558-5646.1991.tb02692.x

Wei W et al. 2022. Rapid evolution of mutation rate and spectrum in response to environmental and population-genetic challenges. Nat Commun. 13(1):4752. 10.1038/s41467-022-32353-6

Weinstein B, Roy SW. 2026. Controlling for life-history traits in vertebrates reveals that effective population size does not affect mutation rate or genome size. Proceedings of the National Academy of Sciences. 123(6):e2519649123. 10.1073/pnas.2519649123

Wilson KP, Blinn DW, Herbst DB. 2001. The Devils Hole Energetics/Community Relationships: Death Valley National Park, California. Two-Year Progress Report to Death Valley National Park.

Wilson MAW, Venditti C, Pagel M, Makova KD. 2011. Do variations in substitution rates and male mutation bias correlate with life-history traits? A study of 32 mammalian genomes. Evol. 65(10):2800–2815. 10.1111/j.1558-5646.2011.01337.x

Wolfe KH, Sharp PM, Li W-H. 1989. Mutation rates differ among regions of the mammalian genome. Nature. 337(6204):283–285. 10.1038/337283a0

Wu FL et al. 2020. A comparison of humans and baboons suggests germline mutation rates do not track cell divisions. PLOS Biology. 18(8):e3000838. 10.1371/journal.pbio.3000838

Xie KT et al. 2019. DNA fragility in the parallel evolution of pelvic reduction in stickleback fish. Science. 363(6422):81–84. 10.1126/science.aan1425

Zhang C et al. 2023. De Novo Mutation Rates in Sticklebacks. Mol Biol Evol. 40(9):msad192. 10.1093/molbev/msad192

Zhang C et al. 2025. Rate of de novo mutations in the three-spined stickleback. Heredity. 134(7):387–395. 10.1038/s41437-025-00767-9

Zhu L, Beichman A, Harris K. 2025. Population size interacts with reproductive longevity to shape the germline mutation rate. Proc Natl Acad Sci U S A. 122(21):e2423311122. 10.1073/pnas.2423311122

