## Supplemental Materials for "Estimate of the mutation rate in the endangered Devils Hole pupfish provides equivocal support for the drift-barrier hypothesis at an outlying extreme"

### Supplemental Figures

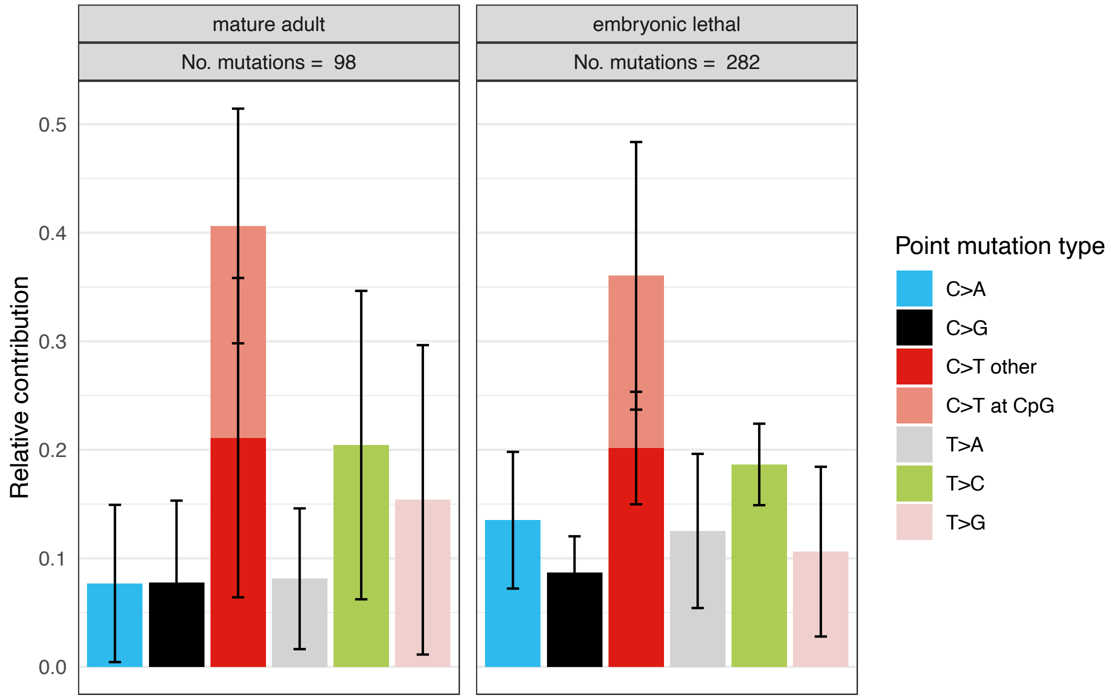


**Supplemental Figure 1.** Single base mutation spectra comparison between *C. diabolis* mature adults and embryonic lethals. Error bars represent 95% confidence interval.

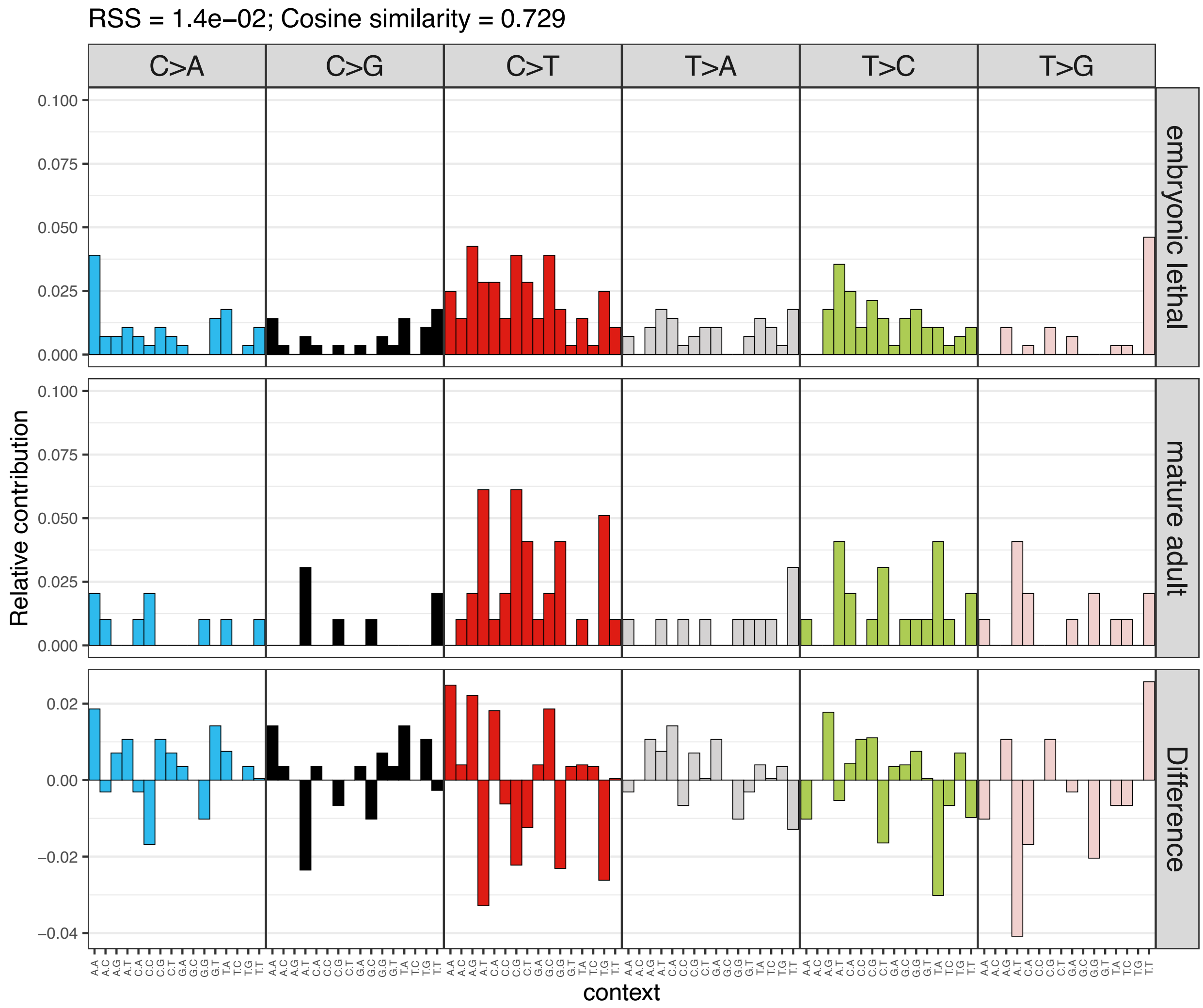


**Supplemental Figure 2.** 96 tri-nucleotide mutation spectra comparison between *C. diabolis* mature adults and embryonic lethals along with difference in relative contribution to each spectra type.

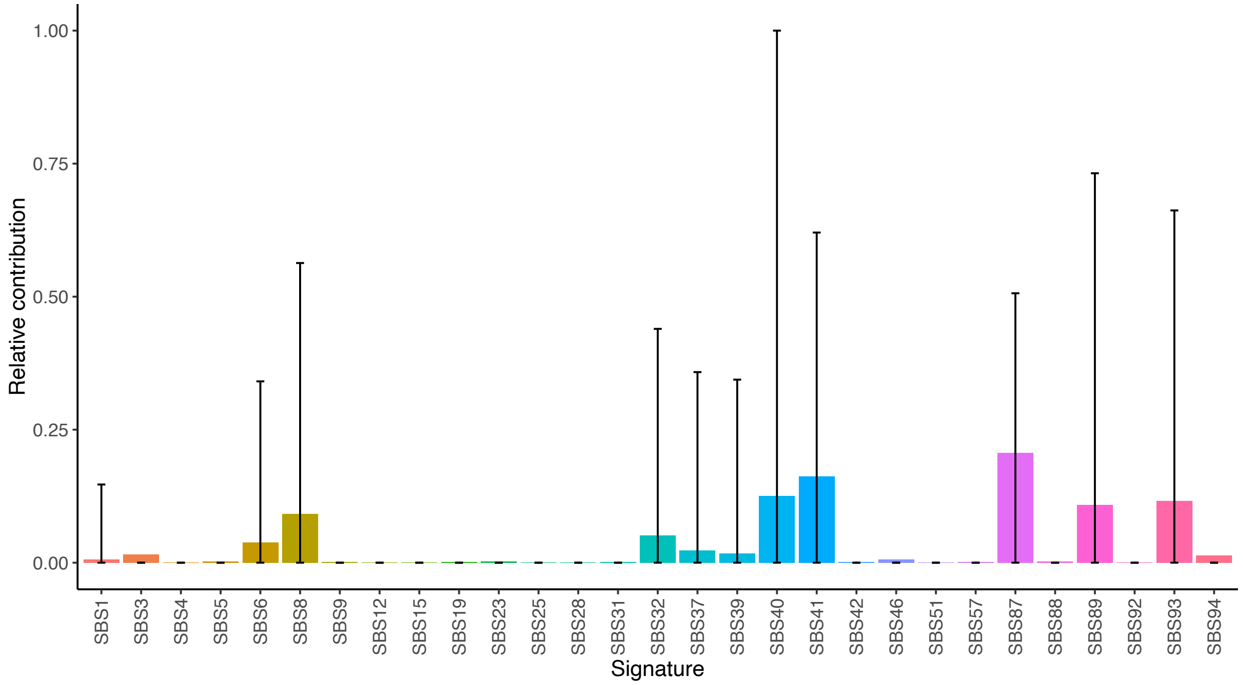


**Supplemental Figure 3.** *C. diabolis* mutational signature and known sequencing artifact signature fitting using MutationalPatterns (Blokzijl et al. 2018). Signatures were refitted with max delta = 0.01 to avoid overfitting and 1,000 bootstraps to assess the signature refitting stability. Error bars represent 95% confidence interval.

### Supplemental Tables

**Supplemental Table 1.** Individual metadata including sample IDs, source, population, sampling year, and sequencing coverage.

| Individual | Source | Population | Year | Coverage group | Coverage |
|---|---|---|---|---|---|
| DH1 | Wild | Mature adult | 2019 | medium | 15.5 |
| DH2 | Wild | Mature adult | 2019 | medium | 18.2 |
| DH3 | Wild | Mature adult | 2019 | medium | 15.5 |
| DH4 | Wild | Mature adult | 2019 | medium | 16.1 |
| DH6 | Wild | Mature adult | 2020 | medium | 16.3 |
| DH7 | Wild | Mature adult | 2020 | medium | 17.1 |
| DH8 | Wild | Mature adult | 2020 | medium | 16.2 |
| DH9 | Wild | Mature adult | 2020 | medium | 15.8 |
| DH10 | Wild | Mature adult | 2020 | medium | 18.1 |
| DH11 | Wild | Mature adult | 2020 | medium | 16.5 |
| DH12 | Wild | Mature adult | 2020 | medium | 16.0 |
| DH13 | Wild | Mature adult | 2020 | medium | 17.2 |
| DH14 | Wild | Mature adult | 2020 | medium | 16.0 |
| DH15 | Wild | Mature adult | 2020 | medium | 20.1 |
| DH16 | Wild | Mature adult | 2020 | medium | 16.6 |
| DH17 | Wild | Embryonic lethal | 2023 | medium | 18.3 |
| DH18 | Wild | Embryonic lethal | 2023 | medium | 24.4 |
| PR-Dtub1 | Captive | Embryonic lethal | 2023 | medium | 18.9 |
| PR-Dtub10 | Captive | Embryonic lethal | 2023 | medium | 21.5 |
| PR-Dtub11 | Captive | Embryonic lethal | 2023 | medium | 19.0 |
| PR-Dtub12 | Captive | Embryonic lethal | 2023 | medium | 16.9 |
| PR-Dtub2 | Captive | Embryonic lethal | 2023 | medium | 18.4 |
| PR-Dtub3 | Captive | Embryonic lethal | 2023 | medium | 22.6 |
| PR-Dtub4 | Captive | Embryonic lethal | 2023 | medium | 16.3 |
| PR-Dtub5 | Captive | Embryonic lethal | 2023 | medium | 18.6 |
| PR-Dtub6 | Captive | Embryonic lethal | 2023 | medium | 21.3 |
| PR-Dtub7 | Captive | Embryonic lethal | 2023 | medium | 16.1 |
| RT1 | Captive | Mature adult | 2020 | medium | 25.1 |
| RT2 | Captive | Mature adult | 2019 | medium | 20.7 |
| RT3 | Captive | Mature adult | 2019 | medium | 15.1 |
| RT4 | Captive | Mature adult | 2019 | medium | 17.3 |
| RT5 | Captive | Mature adult | 2019 | medium | 24.5 |
| RT6 | Captive | Mature adult | 2019 | medium | 23.5 |
| RT7 | Captive | Mature adult | 2019 | medium | 19.2 |
| RT8 | Captive | Mature adult | 2019 | medium | 17.1 |
| RT9 | Captive | Mature adult | 2020 | medium | 16.6 |
| RT10 | Captive | Mature adult | 2020 | high | 112.8 |
| RT11 | Captive | Mature adult | 2020 | high | 87.7 |
| RT12 | Captive | Mature adult | 2019 | medium | 22.3 |

| | | | | | |
|---|---|---|---|---|---|
| RT13 | Captive | Mature adult | 2019 | medium | 17.9 |
| RT14 | Captive | Mature adult | 2019 | high | 94.8 |
| RT15 | Captive | Mature adult | 2019 | high | 16.7 |
| RT16 | Captive | Mature adult | 2020 | medium | 17.8 |
| RT17 | Captive | Mature adult | 2020 | medium | 97.0 |
| RT18 | Captive | Mature adult | 2020 | medium | 15.8 |
| RT19 | Captive | Mature adult | 2020 | medium | 18.9 |
| RT20 | Captive | Mature adult | 2020 | medium | 18.7 |
| RT21 | Captive | Mature adult | 2020 | medium | 17.0 |
| RT22 | Captive | Mature adult | 2020 | medium | 21.9 |
| RT23 | Captive | Mature adult | 2020 | medium | 24.8 |
| RT24 | Captive | Mature adult | 2020 | medium | 15.5 |
| RT25 | Captive | Mature adult | 2020 | medium | 20.9 |
| RT26 | Captive | Embryonic lethal | 2023 | medium | 19.9 |
| RT29 | Captive | Embryonic lethal | 2023 | medium | 24.1 |
| RT31 | Captive | Embryonic lethal | 2023 | medium | 16.7 |
| RT33 | Captive | Embryonic lethal | 2023 | medium | 18.1 |
| RT34 | Captive | Embryonic lethal | 2023 | medium | 17.2 |
| RT35 | Captive | Embryonic lethal | 2023 | medium | 18.1 |
| RT37 | Captive | Embryonic lethal | 2023 | medium | 18.2 |
| RT38 | Captive | Embryonic lethal | 2023 | medium | 16.1 |
| RT39 | Captive | Embryonic lethal | 2023 | medium | 15.2 |
| RT40 | Captive | Embryonic lethal | 2023 | medium | 16.9 |

**Supplemental Table 2.** Mutation data per individual with at least one autozygous segment greater than 5 Mb. The number of heterozygous variants and singletons are restricted to those found within the autozygous segments. The number of callable bp, mean autozygous segment size, and total autozygous account for a mappability mask and 1 Mb removed from both ends of the autozygous segment. The singleton rate was calculated as the number of singletons / (callable bp * 2), weighted by the amount of number of total autozygous bp per individual.

| Individual | Population | Num. segments (> 5 Mb) | Num. het. variants | Num. singletons | Callable bp | Mean seg. size | Total autozygous bp |
|---|---|---|---|---|---|---|---|
| DH10 | adult | 2 | 3 | 0 | 7638523 | 4.22E+06 | 8.45E+06 |
| DH11 | adult | 1 | 2 | 0 | 2791812 | 3402796 | 3.40E+06 |
| DH13 | adult | 1 | 0 | 0 | 4695380 | 5111477 | 5.11E+06 |
| DH14 | adult | 2 | 1 | 0 | 6512353 | 3.69E+06 | 7.37E+06 |
| DH16 | adult | 2 | 0 | 0 | 9368251 | 5.47E+06 | 1.09E+07 |
| DH17 | lethal | 6 | 10 | 5 | 24459591 | 4.68E+06 | 2.81E+07 |
| DH2 | adult | 2 | 5 | 1 | 11364238 | 6253940 | 1.25E+07 |
| DH9 | adult | 5 | 3 | 0 | 26286920 | 5.88E+06 | 2.94E+07 |
| PR-Dtub1 | lethal | 46 | 10 | 0 | 267142843 | 6.60E+06 | 3.03E+08 |
| PR-Dtub11 | lethal | 2 | 1 | 0 | 7066516 | 3856822 | 7.71E+06 |
| PR-Dtub2 | lethal | 17 | 18 | 3 | 106179128 | 7.08E+06 | 1.20E+08 |
| PR-Dtub3 | lethal | 12 | 8 | 1 | 64968315 | 6056092 | 7.27E+07 |
| PR-Dtub4 | lethal | 5 | 5 | 3 | 25717557 | 5.74E+06 | 2.87E+07 |
| PR-Dtub5 | lethal | 15 | 12 | 6 | 83575651 | 6290399 | 9.44E+07 |
| PR-Dtub6 | lethal | 8 | 6 | 0 | 33360202 | 4.70E+06 | 3.76E+07 |
| PR-Dtub7 | lethal | 20 | 16 | 7 | 116026295 | 6.61E+06 | 1.32E+08 |
| RT10 | adult | 29 | 15 | 3 | 164570633 | 6.43E+06 | 1.87E+08 |
| RT11 | adult | 29 | 18 | 1 | 160111060 | 6.32E+06 | 1.83E+08 |
| RT12 | adult | 5 | 0 | 0 | 17594212 | 3.93E+06 | 1.96E+07 |
| RT13 | adult | 2 | 0 | 0 | 5695656 | 3147862 | 6.30E+06 |
| RT14 | adult | 32 | 10 | 2 | 176792118 | 6.29E+06 | 2.01E+08 |
| RT16 | adult | 2 | 1 | 0 | 7339066 | 4061953 | 8.12E+06 |
| RT17 | adult | 28 | 16 | 5 | 162916266 | 6.74E+06 | 1.89E+08 |
| RT19 | adult | 8 | 0 | 0 | 26194127 | 3.85E+06 | 3.08E+07 |
| RT2 | adult | 1 | 2 | 0 | 4318209 | 4826527 | 4.83E+06 |
| RT20 | adult | 5 | 0 | 0 | 15606730 | 3825153 | 1.91E+07 |
| RT21 | adult | 1 | 0 | 0 | 3484446 | 3972832 | 3.97E+06 |
| RT22 | adult | 14 | 6 | 0 | 72144118 | 5855248 | 8.20E+07 |
| RT23 | adult | 4 | 1 | 0 | 14804331 | 4.31E+06 | 1.73E+07 |
| RT26 | lethal | 24 | 29 | 7 | 153286291 | 7.22E+06 | 1.73E+08 |

| | | | | | | | |
|---|---|---|---|---|---|---|---|
| RT31 | lethal | 27 | 12 | 7 | 187258124 | 7.83E+06 | 2.11E+08 |
| RT34 | lethal | 24 | 30 | 5 | 167170282 | 7.84E+06 | 1.88E+08 |
| RT35 | lethal | 27 | 30 | 5 | 174379292 | 7.28E+06 | 1.96E+08 |
| RT37 | lethal | 27 | 29 | 8 | 163605058 | 6.90E+06 | 1.86E+08 |
| RT38 | lethal | 17 | 23 | 5 | 121150667 | 8.06E+06 | 1.37E+08 |
| RT39 | lethal | 19 | 16 | 1 | 124998013 | 7.38E+06 | 1.40E+08 |
| RT4 | adult | 8 | 2 | 1 | 26633819 | 3.83E+06 | 3.06E+07 |
| RT40 | lethal | 30 | 37 | 8 | 184090125 | 6.93E+06 | 2.08E+08 |
| RT5 | adult | 6 | 3 | 1 | 21464585 | 4.10E+06 | 2.46E+07 |
| RT6 | adult | 8 | 0 | 0 | 26270943 | 3.76E+06 | 3.01E+07 |
| RT7 | adult | 2 | 0 | 0 | 6894110 | 4102634 | 8.21E+06 |
| RT8 | adult | 1 | 0 | 0 | 5767023 | 6649120 | 6.65E+06 |
